## Supplementary file 1 for "Antisense oligonucleotide therapy rescues disturbed brain rhythms and sleep in juvenile and adult mouse models of Angelman syndrome"

| **Mouse phenotyping tests** | **Age of *Ube3a* restoration** | | | | | | | | | | | | | | | | |
| --- | --- | --- | --- | --- | --- | --- | --- | --- | --- | --- | --- | --- | --- | --- | --- | --- | --- |
|  | **Embryonic rescue** | | | | **Neonatal rescue** | | | | **Juvenile rescue** | | | **Adult rescue** | | | | | |
|  | E0 (Silva-Santos et al., 2015) | E0 (Rotaru et al., 2018) | E12.5 (Sonzogni et al., 2020) | E15.5 and P1 (Wolter et al., 2020) | P0 (Schmid, et al., 2021) | P1 (Silva-Santos et al., 2015) | P1 (Milazzo et al., 2021) | P1 (Judson et al., 2021) | 3 weeks (Silva-Santos et al., 2015) | 3 weeks (Gu et al., 2018) | 3 weeks (Milazzo et al., 2021) | 6 weeks (Silva-Santos et al., 2015) | 8-18 weeks (Gu et al., 2018) | 14 weeks (Silva-Santos et al., 2015) | 14 weeks (Rotaru et al., 2018) | 2-4 months (Meng et al., 2015) | Adult age  (Daily et al., 2011) |
| Method of restoration | Ube3a^Stop^ and CAG-Cre | Ube3a^Stop^ and CAG-Cre | Ube3a^Stop^ and Nestin-Cre | AAV-Cas9 targeting Ube3a-ATS | AAV-Cas9 targeting Ube3a-ATS | Ube3a^Stop^ and CAG-Cre^ERT^ | ASO targeting Ube3a-ATS | AAV- hUBE3A | Ube3a^Stop^ and CAG-Cre^ERT^ | Ube3a^Stop^ and CAG-Cre^ERT^ | ASO targeting Ube3a-ATS | Ube3a^Stop^ and CAG-Cre^ERT^ | Ube3a^Stop^ and CAG-Cre^ERT^ | Ube3a^Stop^ and CAG-Cre^ERT^ | Ube3a^Stop^ and CAG-Cre^ERT^ | ASO targeting Ube3a-ATS | AAV-mUbe3a |
| Protein level (% of WT) | > 80% | – | ~ 100% | 20-40 % | ~27% | ~ 50% | 40-80 % | ~ 100% | > 70% | ~ 80% | 40-100 % | > 70% | ~ 80% | > 70% | ~ 100% | ~ 40% | ~ 100% |
| Rotarod | Rescued | – | Rescued | Improved | Improved | Rescued | **Failed** | Improved | Rescued | – | – | **Failed** | – | **Failed** | – | **Failed** | – |
| Marble burying | Rescued | – | Rescued | **Failed** | Improved | **Failed** | **Failed** | **Failed** | **Failed** | – | – | **Failed** | – | **Failed** | – | **Failed** | – |
| Open field - travel distance | Rescued | – | – | **Failed** | **Failed** | Rescued | Rescued | **Failed** | **Failed** | – | – | **Failed** | – | **Failed** | – | **Failed** | – |
| Open field - time in center | – | – | – | Rescued | – | – | – | – | – | – | – | **–** | – | – | – | – | – |
| Nest building | Rescued | – | Rescued | – | Improved | **Failed** | **Failed** | Improved | **Failed** | – | – | **Failed** | – | **Failed** | – | – | – |
| Forced swim | Rescued | – | Rescued | – | – | **Failed** | Rescued | – | **Failed** | – | – | **Failed** | – | **Failed** | – | – | – |
| Contextual fear memory | – | – | – | – | – | – | – | – | – | – | – | – | – | – | – | Rescued | Rescued |
| Morris water maze | – | – | – | – | – | – | – | – | – | – | – | – | – | – | – | – | Improved |
| Hindlimb clasping | – | – | – | Rescued | – | – | – | – | – | – | – | – | – | – | – | – | – |
| Body weight (Obesity) | – | – | – | Rescued | Improved | – | – | **Failed** | – | – | – | – | – | – | – | Rescued | – |
| Brain weight | – | – | – | Improved | – | – | Improved | **Failed** | – | – | – | – | – | – | – | – | – |
| Audiogenic seizure | Rescued | – | – | – | – | – | Rescued | Rescued | **Failed** | – | Rescued | – | – | – | – | – | – |
| Flurothyl-induced seizure | – | – | – | – | – | – | – | – | – | Rescued | – | – | **Failed** | – | – | – | – |
| Synaptic plasticity (LTP) | – | – | – | – | – | – | Rescued | – | Rescued | – | Improved | – | – | Rescued | – | – | Rescued |
| Spontaneous inhibitory synaptic inputs | – | Rescued | – | – | – | – | – | – | – | – | – | – | – | – | Rescued | – | – |
| Spontaneous excitatory synaptic inputs | – | Rescued | – | – | – | – | – | – | – | – | – | – | – | – | Rescued | – | – |
| Excitability of fast spiking interneurons | – | – | – | – | – | – | – | – | – | – | – | – | – | – | Rescued | – | – |

**The definition of “rescued”, “improved”, and “failed” in the table:**

Rescued: the difference between treatment and mutant group has *P* < 0.05 AND the difference between treatment and WT group has *P* ≥ 0.05.

Improved: the difference between treatment and mutant group has *P* < 0.05 AND the difference between treatment and WT group has *P* < 0.05.

Failed: the difference between treatment and mutant group has *P* > 0.05.
